# Can In Silico Models Predict Drug-Induced Cardiac Risk in Vulnerable Populations?

**DOI:** 10.1101/2025.09.22.677793

**Authors:** Paula Dominguez-Gomez, Pablo Gonzalez-Martin, Laura Baldo-Canut, Eva Casoni, Ani Amar, Jose M. Pozo, Constantine Butakoff, Mariano Vazquez, Jazmin Aguado-Sierra

**Affiliations:** Elem Biotech, Pier 07, Via Laietana, 26, Barcelona, 08003, Spain; University Pompeu Fabra, Carrer de Tànger, 122-140, Barcelona, 08018, Spain; Barcelona Supercomputing Center, Plaça d’Eusebi Güell, 1-3, Barcelona, 08034, Spain

**Keywords:** cardiac modelling, QT prolongation, arrhythmia, drug safety, cardiomyopathy, sex differences

## Abstract

This study evaluates virtual cardiac populations for preclinical assessment of drug-induced QT interval prolongation and arrhythmic risk. Traditional predictions often rely on small, healthy cohorts, excluding vulnerable populations. Using computational models of realistic heart anatomies and electrophysiology, we generated a virtual cohort of 512 subjects across healthy and diseased hearts (heart failure, dilated and hypertrophic cardiomyopathy, ischaemia, and myocardial infarction). We assessed QT prolongation and arrhythmic events following administration of moxifloxacin (benchmark antibiotic) and contraindicated drugs including quinidine, bepridil, and flecainide.

Patients with heart failure, hypertrophic and dilated cardiomyopathy showed greater QT prolongation to moxifloxacin, unlike ischaemia and myocardial infarction, which resembled healthy subjects. Females exhibited consistently higher QT prolongation than males. Contraindicated drugs markedly increased arrhythmia risk in populations with heart failure, dilated and hypertrophic cardiomyopathy, and ischaemia, frequently leading to lethal arrhythmias such as Torsades des Pointes or ventricular fibrillation, particularly in females.

These findings demonstrate that computational models capture variability in drug response across pathologies and sexes, offering a predictive framework for preclinical safety evaluations and supporting safer, more personalized drug development strategies.

## 1 Introduction

Drug-induced arrhythmias are irregular heart rhythms caused by disruptions in the heart’s electrical activity, with outcomes ranging from non-lethal events, such as benign palpitations or premature ventricular contractions (PVCs), to life-threatening conditions like ventricular fibrillation (VF) or Torsades de Pointes (TdP) (Shah et al., 2005). Fatal arrhythmias, such as VF, TdP, and asystole, can cause sudden cardiac arrest by severely disrupting effective heart function, while non-fatal arrhythmias like ventricular tachycardia (VT) and frequent PVCs, though less dramatic, remain clinically significant due to their potential to progress to more severe conditions. A key biomarker of drug-induced arrhythmic risk is QT interval prolongation, measured on an electrocardiogram (ECG) from the start of the QRS complex to the end of the T wave. QT prolongation is often linked to drug-induced blockade of the hERG potassium channel, which delays cardiac repolarization and increases the risk of arrhythmias (Frommeyer and Eckardt, 2016). Despite substantial progress, the connection between cellular ion channel blockade and QT prolongation at the organ level remains incompletely understood.

Proarrhythmic risk evaluation in clinical and preclinical settings relies on in vitro tests, animal models, and human clinical trials. However, in vitro assays lack systemic complexity (Kofron et al., 2021), and animal studies are not fully translatable to humans (Kofron et al., 2021). Clinical trials of drug cardiac safety typically focus on young, healthy participants, excluding populations with underlying conditions to minimize confounding variables (U.S. Food and Drug Administration, 2005). These exclusions limit the applicability of safety data to diverse patient populations, leaving significant gaps in understanding drug-induced risks for individuals with cardiovascular diseases or other comorbidities (Gross et al., 2022). For instance, patients with heart failure or ischaemia may exhibit altered ion channel functions that heighten their susceptibility to arrhythmias (Gomez et al., 2014), risks that often remain unaddressed during conventional testing.

Computational models of cardiac electrophysiology offer a promising alternative by enabling in silico simulations of drug effects at cellular, tissue, and organ levels. Over time, these models have evolved to incorporate detailed ion channel dynamics and whole-heart behaviour, allowing for more accurate predictions of drug-induced arrhythmias (Mirams et al., 2011; Z. Li et al., 2019; Passini et al., 2017; Abbasi et al., 2017; Passini et al., 2019; Romero et al., 2018; Zemzemi et al., 2013; Hwang et al., 2019; Costabal et al., 2018; Okada et al., 2018; Wilhelms et al., 2012; Cranford et al., 2018; Peirlinck et al., 2021). They complement traditional safety evaluations by providing a scalable, cost-effective, and ethical framework for assessing risks across a wider range of patient profiles. While advancements in disease modelling have enabled detailed simulations of conditions such as myocardial infarction (León et al., 2019; Decker and Rudy, 2010; Arevalo et al., 2016; López-Pérez et al., 2019), ischaemia (Loewe et al., 2018; Salvador et al., 2021; Loidi and Ferrero, 2022; Martinez-Navarro et al., 2019; Martinez-Navarro et al., 2020; Dutta et al., 2017b), heart failure (Gomez et al., 2014; Mora et al., 2017; Shavik et al., 2021), and other cardiomyopathies (Lyon et al., 2018; Passini et al., 2016; Bhattacharya-Ghosh et al., 2014), these efforts rarely address cardiac drug safety directly or consider the variability introduced by comorbidities (TeBay et al., 2022).

Recent studies have begun to address these gaps. For instance, Myklebust et al. (2024) demonstrated that the choice of fibrosis modeling significantly affects the morphology of ventricular arrhythmias in non-ischemic cardiomyopathy, highlighting the importance of accurate structural representations in simulations. Similarly, Coleman et al. (2024) investigated the effects of ranolazine on the arrhythmic substrate in hypertrophic cardiomyopathy, providing insights into the drug’s impact on repolarization and arrhythmic risk. Additionally, Llopis-Lorente et al. (2023) combined pharmacokinetic and electrophysiological models to assess drug-induced arrhythmogenicity, incorporating factors like sex and renal function to predict proarrhythmic outcomes more accurately. These studies underscore the necessity of integrating detailed structural, pharmacological, and physiological data to enhance the predictive power of in silico models.

This gap raises a fundamental question: can in silico models help predict drug-induced arrhythmic risk in clinically vulnerable populations? This study explores that question by leveraging virtual populations to assess proarrhythmic risks across a wide spectrum of patient profiles, including individuals with underlying cardiac pathologies. Using anatomically and electrophysiologically detailed sex-specific computational heart models, we constructed virtual populations representing both healthy and diseased hearts, specifically heart failure, dilated and hypertrophic cardiomyopathy, ischaemia, and myocardial infarction. In total, 512 virtual subjects were generated (32 per anatomy–condition combination), capturing interindividual variability through systematic variation of key ionic conductances. By simulating these pathological states and incorporating interindividual variability, this approach moves beyond conventional safety assessments focused solely on healthy cohorts. It offers a testbed to examine how sex, disease, and physiological context interact with drug effects, aiming to provide earlier and more nuanced insights into arrhythmic risk before such risks emerge in the clinic.

## 2 Methodology

The following section will be divided in three subsections: (i) Virtual Population generation; (ii) Pathological modelling and (iii) the electrophysiological validation strategy employed. A descriptive representation can be found in Figure 1.

**Figure 1:**
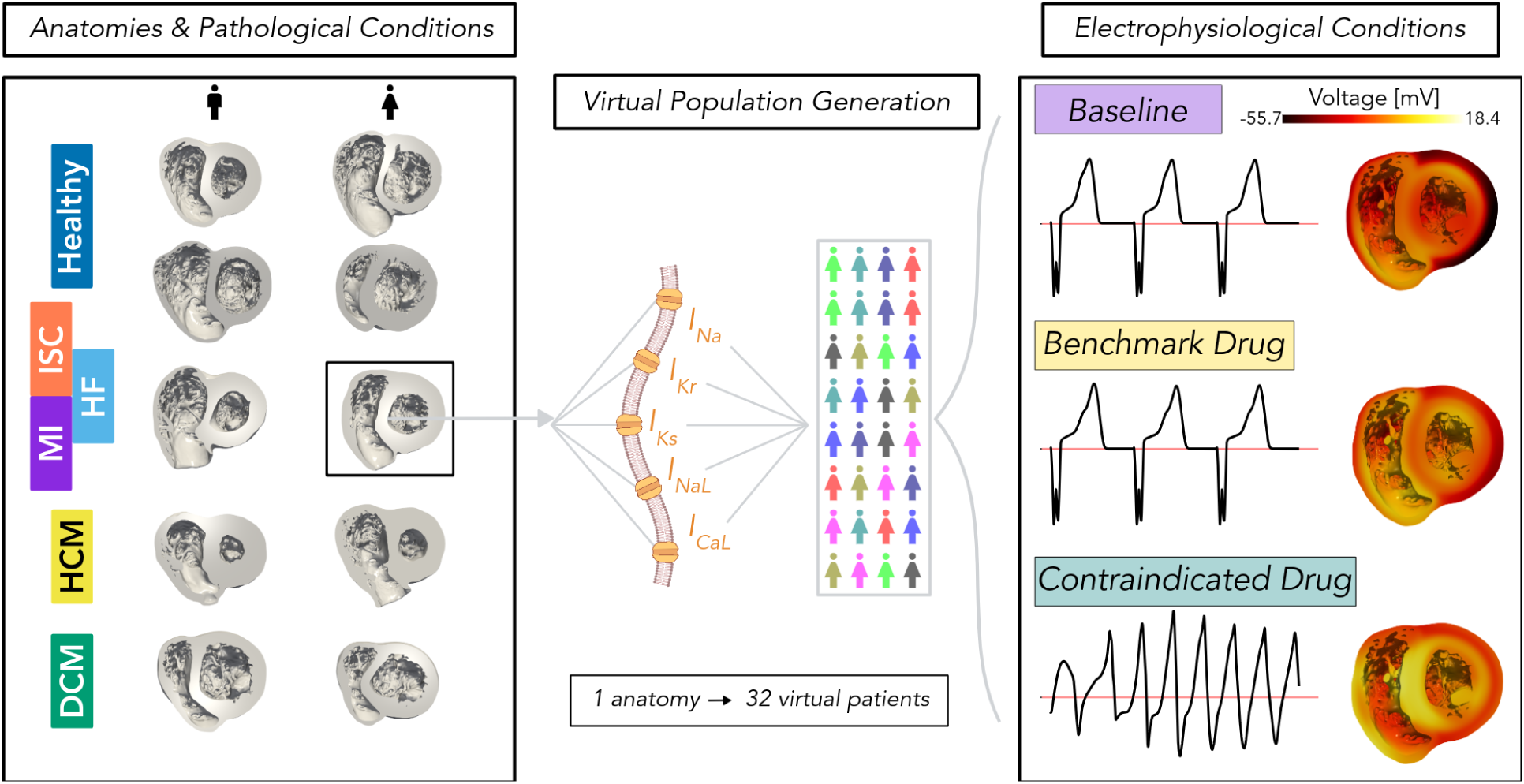
Overview of study methodology. The baseline population captures anatomical variability, including male and female anatomies, as well as specific cardiac pathologies: heart failure (HF), ischaemia (ISC), myocardial infarction (MI), dilated cardiomyopathy (DCM), and hypertrophic cardiomyopathy (HCM). For each combination of anatomy and physiological condition, 32 virtual patients are generated by varying ionic conductances. Each virtual patient is subsequently evaluated under baseline and drug-exposure (benchmark and contraindicated conditions.

### 2.1 Virtual Population Generation

#### 2.1.1 Anatomy Selection

For this study, retrospectively collected data from the Visible Heart Lab library (University of Minnesota Visible Heart Laboratories, 2025) at the University of Minnesota was utilized. Specifically, eight biventricular cardiac geometries (see Table 1 and Figure 2) were included, reconstructed from high-resolution magnetic resonance imaging (MRI) data. The selection aimed to capture a wide spectrum of anatomical variability, including differences in sex, body mass index (BMI), and age, to ensure a representative set of morphologies. Additionally, to address the limited availability of clinical datasets for this study, two artificially dilated biventricular cardiac geometries were created by transforming detailed heart models of each sex to match the size of a reference heart with dilated cardiomyopathy. For further details see Appendix B.

**Figure 2:**
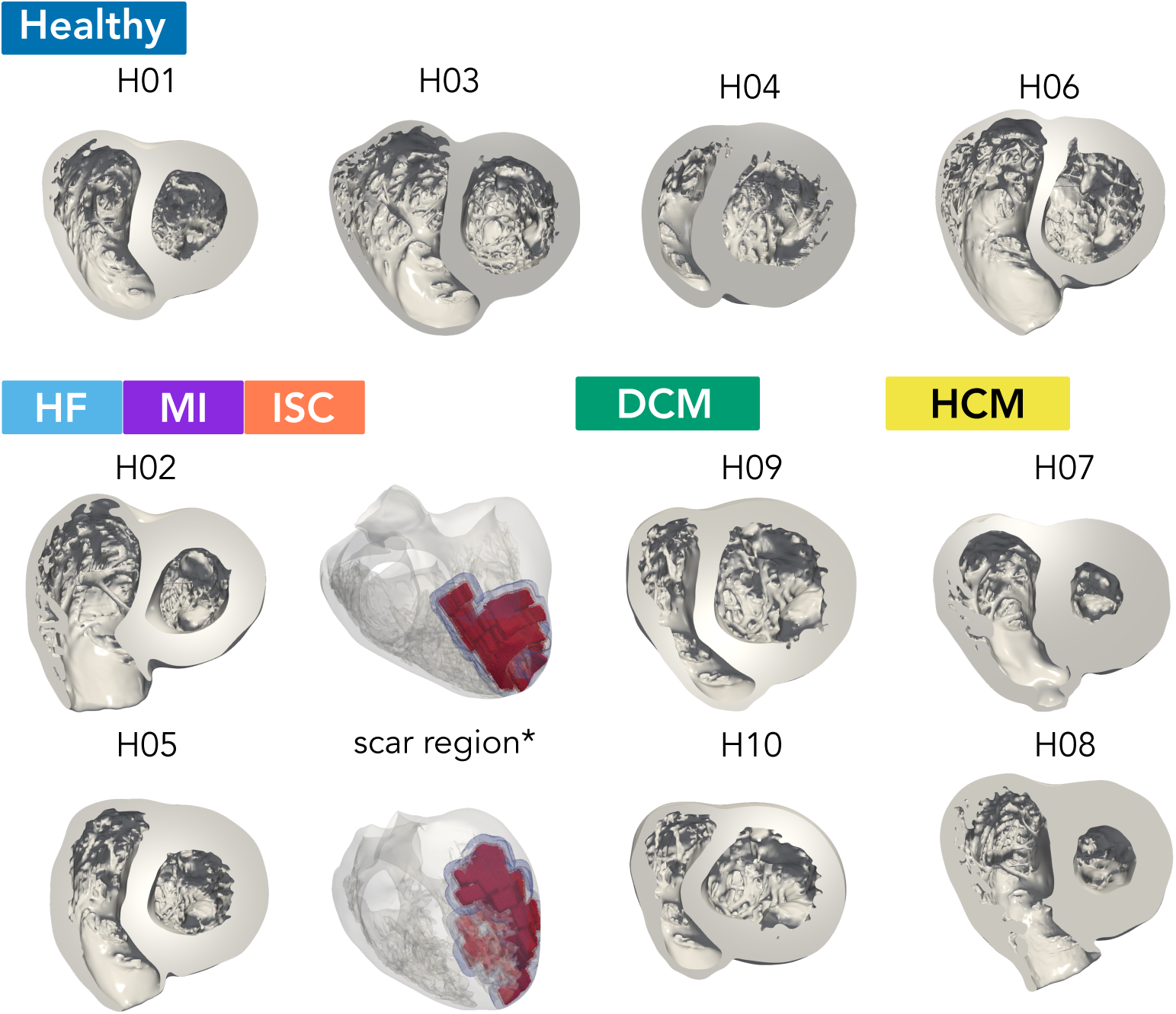
Reconstructed hearts from the Visible Heart Lab library, including AG hearts. Hearts H02 and H05 were both used for healthy, HF, ISC and MI conditions. ^∗^Scars were included just for ISC and MI conditions.

**Table 1:** Selected hearts from the Visible Heart Lab library, including AG (artificially generated) hearts. Myocardial diffusion in fibre direction indicates the values adjusted to achieve appropriate QT and QRS measurements for the corresponding age and sex group according to reference values (Mason et al., 2007).

| Heart ID | H01 | H02 | H03 | H04 | H05 | H06 | H07 | H08 | H09 | H10 |
| --- | --- | --- | --- | --- | --- | --- | --- | --- | --- | --- |
| Sex | M | M | M | F | F | F | M | F | M | F |
| Age | 36 | 61 | 23 | 58 | 58 | 24 | 84 | 78 | AG | AG |
| BMI | 33.5 | 21.8 | 26.2 | 31.7 | 22.0 | 31.2 | 30.2 | 25.8 | AG | AG |
| Volume (cm<sup>3</sup>) | 352 | 266 | 317 | 455 | 257 | 217 | 319 | 284 | 519 | 324 |
| % of trabeculations | 15 | 19 | 23 | 19 | 13 | 22 | 10 | 12 | 32 | 35 |
| Cardiac clinical history | - | - | - | - | - | - | HCM | HCM | DCM | DCM |
| LV mean wall thickness (mm) | 13.80 | 11.50 | 9.151 | 12.09 | 12.53 | 8.585 | 19.28 | 19.72 | 10.04 | 8.091 |
| Myocardial diffusion in fibre direction (m<sup>2</sup>/ms) | 0.00450 | 0.00400 | 0.00470 | 0.00450 | 0.00375 | 0.00455 | 0.00290 | 0.00350 | 0.00235 | 0.00235 |

The eight anatomies were then used to generate the healthy virtual population (H01–H06) and the pathological cohorts as follows: HF, MI, and ISC populations were generated using again H02 and H05; HCM models were derived from H07 and H08; and DCM models were based on the dilated anatomies H09 and H10.

#### 2.1.2 Anatomy reconstruction and meshing

The anatomical reconstruction of the hearts followed the same methodology expressed in Gonzalez-Martin et al. (2023). The open-source software *3D Slicer* (Version 5.2.1) (Kikinis et al., 2013) was used. Given the high resolution of ex vivo heart images, thresholding based on Renyi entropy (Beare, 2011) was applied, to achieve a preliminary segmentation of cardiac tissue. The segmentation was then refined by leaving only the segmentation of the ventricular myocardium and trabeculated tissue. With the mask defined, a marching cubes algorithm (University of Utah, 2025) was applied to reconstruct the detailed 3D heart. Finally, post-processing was performed using MeshMixer (Version 3.5.4) (Schmidt and Singh, 2010) to remove and/or correct artefacts from segmentation and reconstruction. Lastly, the final reconstructions were meshed using ANSA (Systems, 2025) aiming to maintain edge lengths at approximately 328 *µ*m. Final meshes averaged from 15 to 100 million tetrahedral elements. Reconstructions can be found in Figure 2.

#### 2.1.3 Electrophysiological Modelling

Cardiac electrophysiology, finite element modelling was solved using Alya (Vázquez et al., 2016; Vázquez et al., 2011). Electrophysiology was modelled using the monodomain approximation coupled to the O’Hara-Rudy (ORd) cellular model (O’Hara et al., 2011), using a modified version of the ORd model by Passini et al. (2016) with modified conductances as described in Dutta et al. (2017a). Sex-specific ion channel expression was applied as described by Iseppe et al. (2021), which optimises Yang and Clancy (2012) approach. Sex was assigned based on the original MRI-derived heart geometries, resulting in sex differences being represented at both anatomical and electrophysiological levels. This parametrization was used as the reference cellular model in this study (Table 2).

**Table 2:**
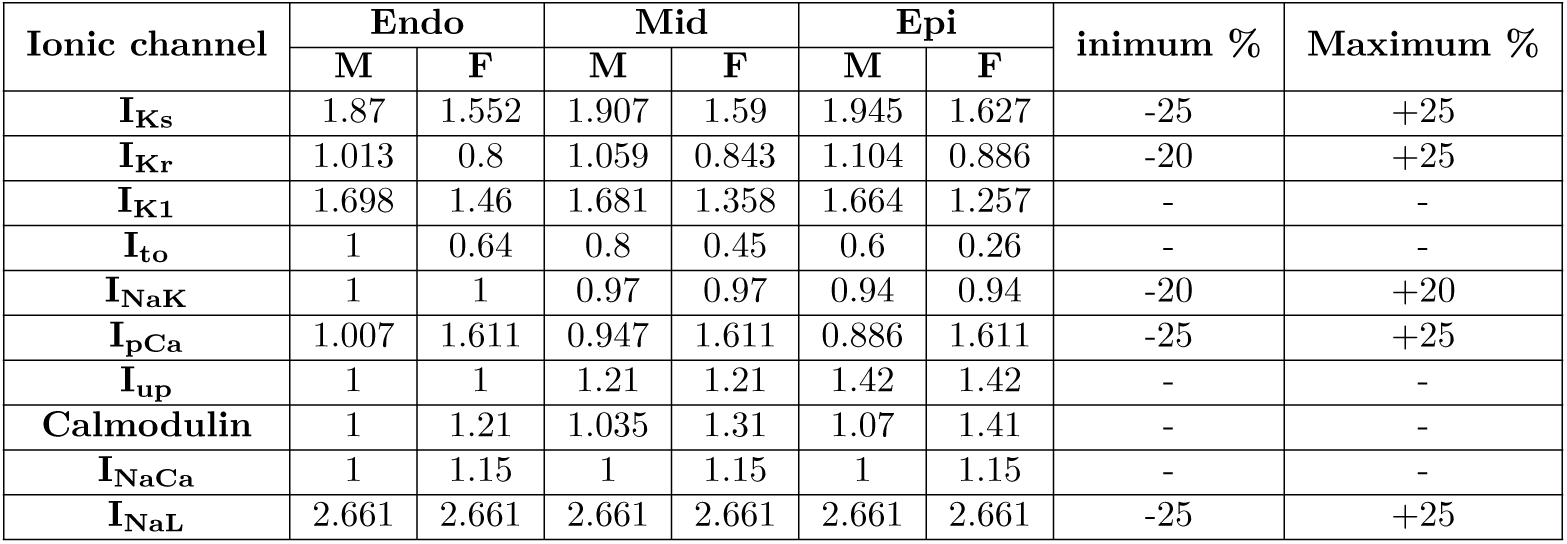
Differences in ionic channel conductance scaling factors by sex, based on the ORd model, in order to reproduce the reference cellular model (Passini et al., 2016; Iseppe et al., 2021). Percentage adjustments made to account for variability in the virtual population included.

Electrical activation was initiated following Durrer et al. (1970) early activation points, with 5 mA/cm^3^ applied for 5 ms at specific sites in the biventricular cavities. Specifically, activation sites included two areas on the right ventricular (RV) wall near the insertion of the anterior papillary muscle, a high anterior paraseptal region of the left ventricle (LV) below the mitral valve, a central area on the left surface of the septum, and a posterior LV paraseptal region located at 1*/*3 of the apex-base distance. Cardiac fibres were assigned using a rule-based approach suited for detailed geometries, including trabeculae and papillary muscles (Doste et al., 2019). A detailed explanation is available in Gonzalez-Martin et al. (2023).

To capture cellular heterogeneity, the myocardium was divided into endocardial (33%), mid-myocardial (33%), and epicardial (33%) layers, each with distinct electrophysiological properties. For mid-myocardial cells, in the absence of sex-specific data in the selected cellular model, the modifications applied to endocardial and epicardial cells were averaged.

The inclusion of mid-myocardial cells is based on experimental evidence demonstrating their distinct ion channel expression and repolarization properties in isolated myocytes (Wilson et al., 2011; Liu et al., 1993). While there is ongoing debate regarding their functional role in healthy ventricular myocardium, different studies have confirmed their contribution to transmural repolarization gradients, particularly under disease conditions (Glukhov et al., 2010; Antzelevitch et al., 1999). Given that mid-myocardial cells play a role in arrhythmic dynamics and have been widely employed in full-ventricle electrophysiological simulations, their inclusion ensures a consistent and physiologically plausible framework for our study.

In addition, to approximate Purkinje fibre conduction, an additional endocardial layer with three times faster diffusion properties was incorporated, enabling an efficient representation of rapid ventricular activation.

#### 2.1.4 Virtual Population

A virtual population was generated following the approach described by Muszkiewicz et al. (2016), which has been tested in previous work (Aguado-Sierra et al., 2024b; Aguado-Sierra et al., 2024a) demonstrating its suitability for assessing QT prolongation risk. In this method, variability was introduced in the five ionic channels most influential in action potential duration (*I*_Na_, *I*_Kr_, *I*_Ks_, *I*_CaL_, and *I*_NaL_), with conductance variations constrained to ±20-25% (see Table 2), a range consistent with normal human variability as reported by several studies (Walmsley et al., 2013; Romero et al., 2009; Fink et al., 2008; G.-R. Li et al., 1999; Szentandrassy et al., 2005; Verkerk et al., 2005; Virág et al., 2001).

Due to the limited availability of experimental data on sex-specific ionic current variability, the same range of variation was applied to both male and female phenotypes. For each anatomical model, 32 subjects of the corresponding sex were generated by combinatorially pairing the minimum and maximum values of these five conductances. This approach ensured full coverage of the variability range while maintaining a physiologically plausible, healthy adult cohort, as typically recruited in cardiac safety clinical trials.

### 2.2 Pathological Modelling

#### 2.2.1 Heart Failure

HF is the final stage of many cardiovascular diseases and is characterised by impaired contractile function and increased arrhythmic susceptibility. At the cellular level, HF is marked by action potential prolongation, ion channel remodelling, and alterations in calcium handling, all contributing to the pathological electrophysiological state. The cellular electrophysiological activity of HF was simulated following the approach described by Gomez et al. (2014). This involved applying scaling factors to the reference model parameters, as detailed in Table 3, alongside halving the diffusion values. These modifications were carefully chosen to align with experimental findings, ensuring the model’s biological relevance. To avoid excessively pathological manifestations, the changes implemented from Gomez et al. were reduced by 20%. The male heart H02 and female H05 from Table 1 were selected to create this virtual population, as they represent individuals with average characteristics.

**Table 3:** HF ion channel remodelling (expressed as percentage of change) from Gomez et al. (2014), applied to the reference model.

| Reference model parameter | Regulation in HF remodelling |  |  |
| --- | --- | --- | --- |
|  | Endo | Mid | Epi |
| $I_{NaL}$ | | +80 | |
| $\tau_{hL}$ | | +80 | |
| $I_{to}$ | | -60 | |
| $I_{K1}$ | | -32 | |
| $I_{NaK}$ | | -30 | |
| $I_{CaMKa}$ | | +50 | |
| $J_{leak}$ | | +30 | |
| $K_{rel,Ca}$ | | -20 | |
| $I_{NCX}$ | +60 | +60 | +100 |
| $J_{SERCA}$ | -55 | -40 | -25 |

#### 2.2.2 Dilated Cardiomyopathy

DCM is one of the most common causes of HF and is characterised by heart enlargement with wall thinning and weakened contractile function (Seferović et al., 2019). Despite the different underlying causes, both HF and DCM lead to a reduced ability of the heart to supply oxygenated blood, resulting in similar clinical manifestations and arrhythmic risks. To simulate the effects of DCM, we adapted the HF model by applying it to artificially dilated hearts H09 and H10, described in Table 1. Therefore, using a single computational model to simulate both HF and DCM represents an efficient strategy, as conditions share overlapping patho-physiological features. In particular, the disrupted electrical function of the heart cells can be represented through a common underlying mechanism. However, it should be noted that this model does not account for genetic causes of DCM, which are based on mechanical alterations; such electromechanical modelling is beyond the scope of this study.

#### 2.2.3 Hypertrophic Cardiomyopathy

HCM is a complex cardiomyopathy associated with various abnormalities, including ventricular wall thickening, cardiomyocyte disarray, microvascular ischaemia, fibrosis, and ion channel remodelling, all contributing to heightened arrhythmic risk. Although sarcomeric mutations are common causes, other structural and functional abnormalities significantly impact disease manifestation. To model this pathology, one male and one female heart with HCM (H07 and H08 from Table 1) were selected, featuring left ventricular wall thicknesses of approximately 19.28 mm and 19.72 mm, respectively. Fibre orientation was modelled as normal, except in hypertrophic regions where a 50% random misalignment was applied (Figure 3). Fibrosis was simulated by reducing diffusion by 50% in hypertrophic areas. Ion channel remodelling was implemented using conductance changes from Coppini et al. (2013) and updated by Passini et al. (2016), with modifications reduced by 20% to avoid excessively pathological phenotypes inconsistent with clinical QT interval observations (Table 4).

**Figure 3:**
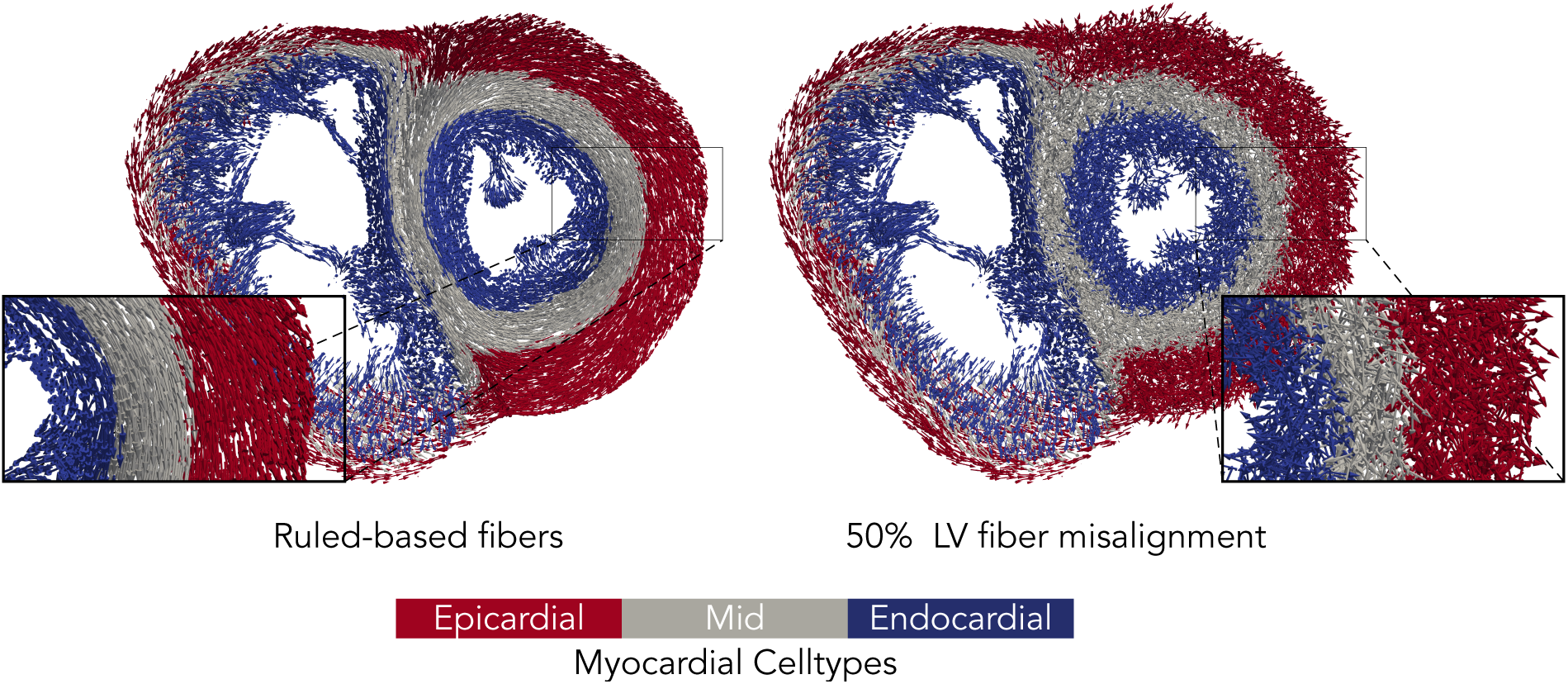
Ruled-based fibres short axis slice of a HCM heart. On the left, original fibre distribution. On the right, 50% fibre misalignment.

**Table 4:** HCM ion channel remodelling (expressed as percentage of change) from Coppini et al. (2013) and Passini et al. (2016), applied to the reference model.

| Reference model parameter | Regulation in HCM remodelling |
| --- | --- |
| $I_{NaL}$ | +107 |
| $I_{to}$ | -85 |
| $I_{CaL}$ | +19 |
| $I_{Kr}$ | -34 |
| $I_{Ks}$ | -27 |
| $I_{K1}$ | -15 |
| $I_{NaCa,i}$ | +34 |
| $I_{NaCa,ss}$ | +34 |
| $I_{NaK}$ | -30 |
| $J_{rel, NP, inf}$ | -20 |
| $J_{rel, CaMK, inf}$ | -20 |
| $J_{up, NP}$ | -43 |
| $J_{up, CaMK}$ | -43 |
| $I_{nab}$ | +107 |
| $\tau I_{CaL}$ fast inactivation | +35 |
| $\tau I_{CaL}$ slow inactivation | +20 |

#### 2.2.4 Ischaemia

ISC arises from reduced blood flow due to coronary artery disease, depriving the heart of oxygen and leading to structural and functional remodelling. Remodelling predominantly affects the core of the ischaemic region and the adjacent border zone (BZ). Following Martinez-Navarro et al. (2020), a 5 mm BZ was defined, with conductances and ionic concentrations values varying smoothly from healthy tissue to full remodelled in the core ischaemic region, as specified in Table 5. This BZ is illustrated in Figure 2. In this work, the spatial location of the ischaemic region was based on MRI segmentations of scars from acutely infarcted human hearts. For the female model (H05 in Table 1), a subendocardial scar from obstruction of the left anterior descending artery was selected. For the male model (H02), a transmural scar from obstruction of the right coronary artery was used, reflecting some common sources of myocardial infarctions.

**Table 5:** ISC extracellular potassium concentration (expressed in mM) and conductance channel remodelling (expressed as percentage of change) from Martinez-Navarro et al. (2020), applied to the reference model.

| Reference model parameter | Regulation in ISC remodelling |
| --- | --- |
| $[K^+]_o$ | 9.5 |
| $I_{K(ATP)}$ | +7 |
| $I_{Na}$ | -25 |
| $I_{CaL}$ | -25 |

#### 2.2.5 Myocardial Infarction

MI is an ischaemic heart disease marked by myocardial necrosis due to coronary artery occlusion, leading to scar formation and remodelling that alters electrical and mechanical properties. Three regions were differentiated in the model: dense scar, heterogeneous scar, and non-scarred tissue, following previously described methodologies (León et al., 2019).

The dense scar core, which lacks viable active myocytes, was modelled as only 10% of the normal extracellular diffusion (Niederer et al., 2011) to ensure existence and continuity of the myocardial tissue. The heterogeneous border zone, composed of viable myocytes and collagen, was modelled with 90% reduced transverse fibre direction diffusion and adjusted ion channel expression levels as detailed in Table 6, based on Decker and Rudy’s cellular model for myocardial infarction (Decker and Rudy, 2010). The diffusion along the longitudinal direction was assumed to remain unaffected, as longitudinal conduction is typically preserved despite changes in the border zone (Arevalo et al., 2016; Yao et al., 2003). Electrical remodelling in the non-infarcted myocardium was characterised using experimental porcine data from invasive electrophysiological studies and cardiac MRI. An S1-S2 restitution protocol was employed, both experimentally and computationally to approximate simulated to experimental restitution curves of the non-scarred post MI tissue. Ion channel conductances in the O’Hara-Rudy model were scaled to account for the remodelling observed. The defined ionic changes for the right and left ventricles are detailed in Table 6.

**Table 6:** MI ion channel conductance scaling factors applied to the reference model, with parameter adjustments specified for the non-infarcted tissue in the left and right ventricles (fitted from experimental data) and the heterogeneous scar tissue (from Decker and Rudy (2010)).

| Reference Model<br>Parameter | Endo |  | Mid |  | Epi |  |
| --- | --- | --- | --- | --- | --- | --- |
|  | M | F | M | F | M | F |
| **Non-Infarcted Tissue (Left Ventricle)** |  |  |  |  |  |  |
| $I_{Na}$ | 0.683 | | 0.266 | | 0.152 | |
| $I_{NaL}$ | 1.863 | | 1.896 | | 1.929 | |
| $I_{CaL}$ | 0.045 | 0.073 | 0.072 | 0.123 | 0.096 | 0.174 |
| $I_{Kr}$ | 0.712 | 0.562 | 0.745 | 0.593 | 0.778 | 0.625 |
| $I_{Ks}$ | 0.453 | 0.376 | 1.967 | 1.639 | 3.540 | 2.961 |
| **Non-Infarcted Tissue (Right Ventricle)** |  |  |  |  |  |  |
| $I_{Na}$ | 0.263 | | 0.535 | | 1.333 | |
| $I_{NaL}$ | 0.958 | | 1.114 | | 3.185 | |
| $I_{CaL}$ | 0.660 | 1.055 | 0.656 | 1.117 | 0.649 | 1.179 |
| $I_{Kr}$ | 1.355 | 1.070 | 1.211 | 0.965 | 1.049 | 0.842 |
| $I_{Ks}$ | 0.547 | 0.454 | 0.808 | 0.674 | 1.079 | 0.903 |
| **Heterogeneous Scar Tissue** |  |  |  |  |  |  |
| $I_{Na}$ | 0.620 | | 0.620 | | 0.620 | |
| $I_{NaL}$ | 2.661 | | 2.661 | | 2.661 | |
| $I_{CaL}$ | 0.695 | 1.112 | 0.653 | 1.112 | 0.611 | 1.112 |
| $I_{Kr}$ | 0.709 | 0.560 | 0.741 | 0.590 | 0.773 | 0.620 |
| $I_{Ks}$ | 1.496 | 1.242 | 1.526 | 1.272 | 1.556 | 1.302 |

Two heart anatomies were selected for this study: female H05 and male H02, as detailed in Table 1. The infarcted scars were selected using the same criteria as for the ischaemic scars.

### 2.3 Modelling drugs

We employed the methodology outlined by Mirams et al. (2011) to account for the effects of drugs on ion channel conductances, utilizing a multi-channel conductance-block formulation as in our previous work (Aguado-Sierra et al., 2024b). The ion channel conductance block is then applied to the conductance *g*_x_ of any of the following ionic channels: *I*_CaL_, *I*_NaL_, *I*_to_, *I*_Ks_, *I*_K1_, *I*_Na_, and *I*_Kr_ (Crumb et al., 2016). We defined the ion channel conductance after drug administration using the following Hill model:

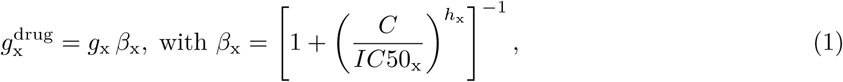

where x represents the specific channel type. In these equations, *g*^drug^ is the conductance of the x-th channel after drug administration, *C* is the drug concentration, *IC*50_x_ is the concentration required to achieve a 50% current blockade of the x-th channel, and *h*_x_ is the corresponding Hill exponent. The blockade *β*_x_ of channel x is characterised by the Hill parameters *h*_x_, the *IC*50_x_, and the drug concentration *C*. It is important to note that *C* is defined as the concentration of the drug in the plasma that is not bound to plasma proteins and is therefore available to exert a pharmacological effect. This is expressed as:

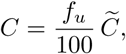

where *C* is the total concentration of the drug in the plasma and *f_u_* (expressed as a percentage) is the free fraction of the drug, representing the ratio of the unbound drug concentration to the total drug concentration.

### 2.4 Drug safety biomarkers

Pseudo-electrocardiograms (pseudo-ECG) were generated by placing each biventricular model within a generic torso constructed from a patient’s computed tomography scan. The cardiac potential was recorded at specific locations to compute the standard 12-lead ECG. Additional details on the pseudo-ECG computation can be found in Gonzalez-Martin et al. (2023).

The pseudo-ECG was used to obtain biomarkers, including the QT interval duration at baseline (QT^bsl^) and after drug administration (QT^drug^). These calculations were performed using an automatic algorithm, as described in Aguado-Sierra et al. (2024b). QT prolongation was defined as:

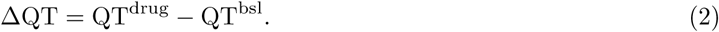

All simulations were done at a constant heart rate of 60 bpm. This ensures that the observed ΔQT effects reflect solely the drug’s influence, facilitating clearer comparisons to clinical data (Garnett et al., 2018).

### 2.5 Electrophysiological Response Validation in Normal and Pathological Cohorts

To validate the pseudo-ECG and associated biomarkers, specifically the QT and QRS intervals derived from our population models under baseline conditions (60 bpm), we conducted a comparative analysis against established normative data. Following the work of Mason et al. (2007), we calibrated our virtual populations to align with reported mean QT values across sex and age groups. Given that clinical studies typically apply correction formulas to account for heart rate variability, we referenced the heart rate-corrected QT interval ranges for 60 bpm: QTc-Bazett, 361 to 457 milliseconds, and QTc-Fridericia, 359 to 445 milliseconds. While some simulated QT values may extend beyond these ranges, they remain within the broader borderline clinical range (approximately 300-500 ms) (Viskin, 2009), where these QT values have still been observed in healthy cohorts.

Drug administration was then tailored to each cohort. All healthy and pathological populations received moxifloxacin (Drugs.com, 2024b) as a benchmark drug to reproduce clinically reported QT prolongation and to compare responses across conditions. Each pathological cohort was additionally exposed to a contraindicated drug selected for its known proarrhythmic risk in that specific condition. This strategy was particularly valuable given the limited clinical data available for diseased populations, enabling the observation of expected proarrhythmic outcomes and validation of model behaviour under clinically relevant scenarios. Input data for the drugs of study can be found in Table 7.

**Table 7:** Selected doses (*C*_1_ and *C*_2_), free fraction (*f_u_*), and ionic channel blocking profiles for the studied drugs. The IC50 and Hill coefficient values were sourced from Crumb et al. (2016).

| Drug | | $\tilde{C}_1$<br>(ng/mL) | $\tilde{C}_2$<br>(ng/mL) | $f_u$<br>(%) | $I_{CaL}$ | $I_{Kr}$ | $I_{K1}$ | $I_{to}$ | $I_{Ks}$ | $I_{NaL}$ | $I_{Na}$ |
| --- | --- | --- | --- | --- | --- | --- | --- | --- | --- | --- | --- |
| Moxifloxacin | IC50 | 1929 | 4663 | 65 | - | 93041 | - | - | 50321 | 382337 | - |
| | $h$ | | | | - | 0.6 | - | - | 1.0 | 1.1 | - |
| Bepridil | IC50 | 1500 | 4500 | 1 | 2808 | 149 | - | - | - | 1814 | 2929 |
| | $h$ | | | | 0.6 | 0.9 | - | - | - | 1.4 | 1.2 |
| Flecainide | IC50 | 200 | 1000 | 55 | 25599 | 692 | - | 9266 | - | 18870 | 6677 |
| | $h$ | | | | 1.4 | 0.8 | - | 0.7 | - | 0.6 | 1.9 |
| Quinidine | IC50 | 400 | 1700 | 16 | - | 343 | - | 3487 | 4899 | - | - |
| | $h$ | | | | - | 1.0 | - | 1.3 | 1.4 | - | - |

Quinidine (Drugs.com, 2024c), an antiarrhythmic that slows heart rate and prolongs the QT interval, was administered to HF patients due to its contraindication in this population because it can worsen HF and increase the risk of proarrhythmic events (Gjesing et al., 2015; Selzer and Wray, 1964; Keren et al., 1981; Sclarovksy et al., 1979). Bepridil (RxList.com, 2022), a calcium channel blocker with strong QT-prolonging and arrhythmogenic properties, was used for DCM, HCM, and MI (Yasuda et al., 2006). For ISC, flecainide (Drugs.com, 2024a), another antiarrhythmic with significant QT-prolonging effects, was chosen due to its contraindication for use in patients with structural heart disease or ischemic conditions due to its proarrhythmic potential (Basza et al., 2023; Winkelmann and Leinberger, 1987).

### 2.6 Computational resources

A total of 3456 simulations were executed, requiring approximately 16 million core hours in total. Each simulation employed 672 computing cores and ran for an average of 8 hours of wall-clock time. These resources enabled high-resolution biventricular electrophysiology simulations across all virtual subjects and conditions.

## 3 Results

### 3.1 Electrophysiological Models’ Evaluation

To evaluate the healthy and diseased phenotypes, action potentials (APs) were analyzed across all modeled conditions (see Figure 4) for the same virtual subject. Healthy AP traces use the ionic scaling factors reported in Table 2 to account for the reference cellular model, whereas pathological traces apply the corresponding remodelling parameters from Tables 3 to 6 over the reference cellular model. In the healthy population, AP durations are within normal ranges (Feher, 2012), with females exhibiting slightly longer durations than males. In HF and DCM, AP durations are markedly prolonged, particularly in endocardial and epicardial cells of females, while male cells display more pronounced mid-myocardial prolongation. These findings are consistent with experimental observations (Lou et al., 2012; Fu and Laurita, 2018; Mages et al., 2021). In HCM, AP durations are elevated across all cell types in both sexes. Notably, the female mid-myocardial cells demonstrate signs of arrhythmogenic behavior in the simulations, reflecting clinical reports of prolonged repolarization and increased arrhythmia susceptibility in HCM patients (Coppini et al., 2013). With respect to ISC and MI, the AP responses correspond specifically to the ischaemic region and heterogeneous scar, respectively, as they are pathologies with local manifestations. Under ISC conditions, AP durations are slightly reduced, in agreement with experimental studies on non-early ischaemic stages (Carmeliet, 1999). In MI, Figure 4 highlights increased generalized AP durations, which aligns with reported experimental data (Song et al., 2024).

**Figure 4:**
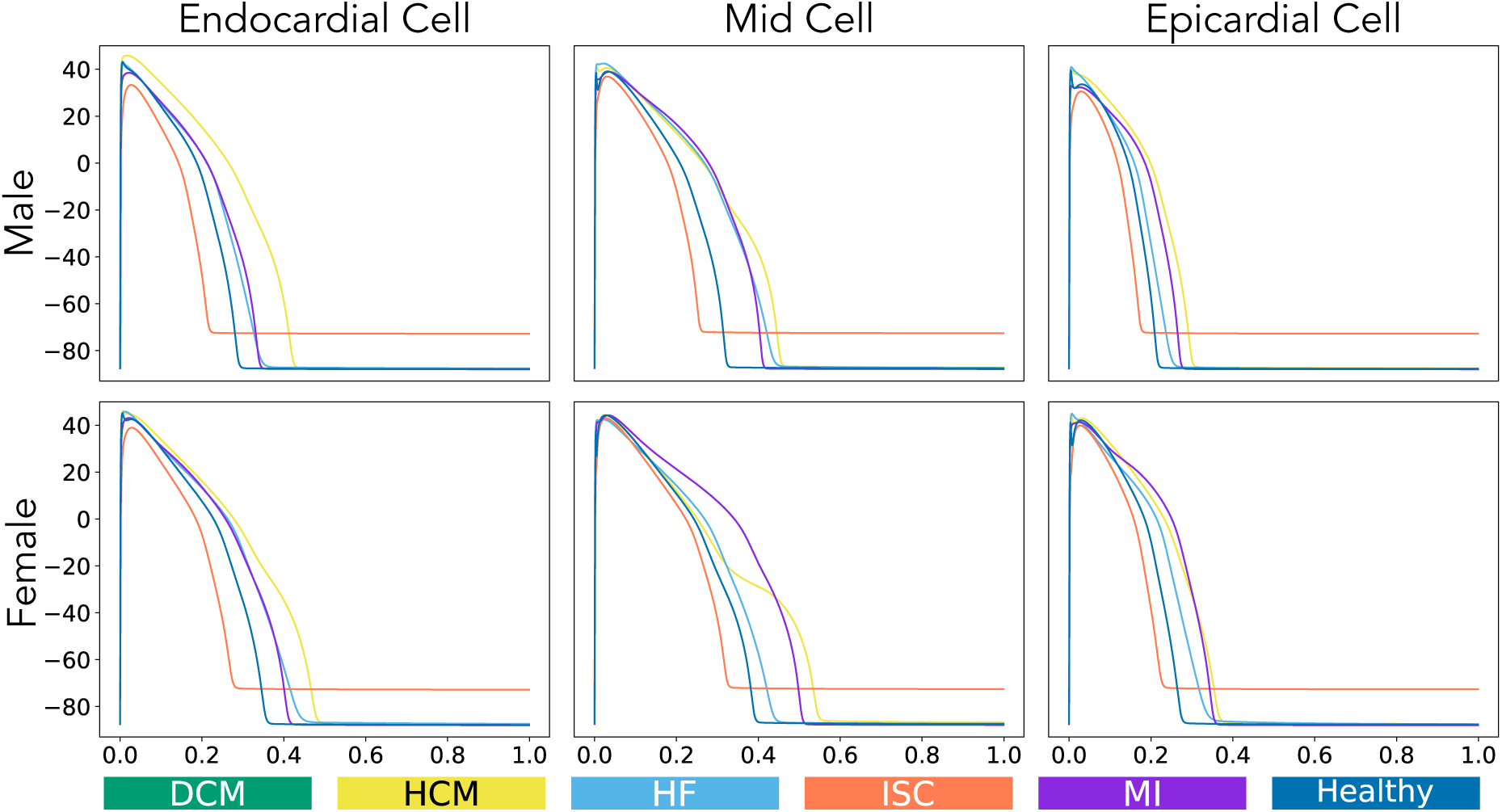
Action potentials of the same single representative virtual subject from different populations under baseline conditions.

At the organ level, these cellular electrical changes are reflected in the ECGs, which provide a macroscopic view of cardiac electrical activity (Figures 5A, 5B). In HF, there is a marked reduction in amplitude and a prolonged duration of QT and QRS complexes compared to healthy individuals, consistent with impaired ventricular function (Miró et al., 2023; Kashani and Barold, 2005). Similarly, in DCM, QT and QRS intervals are prolonged, with generalized voltage reductions aligning with previous studies (Finocchiaro et al., 2020; Alonso et al., 2005; Berger et al., 1997).

**Figure 5:**
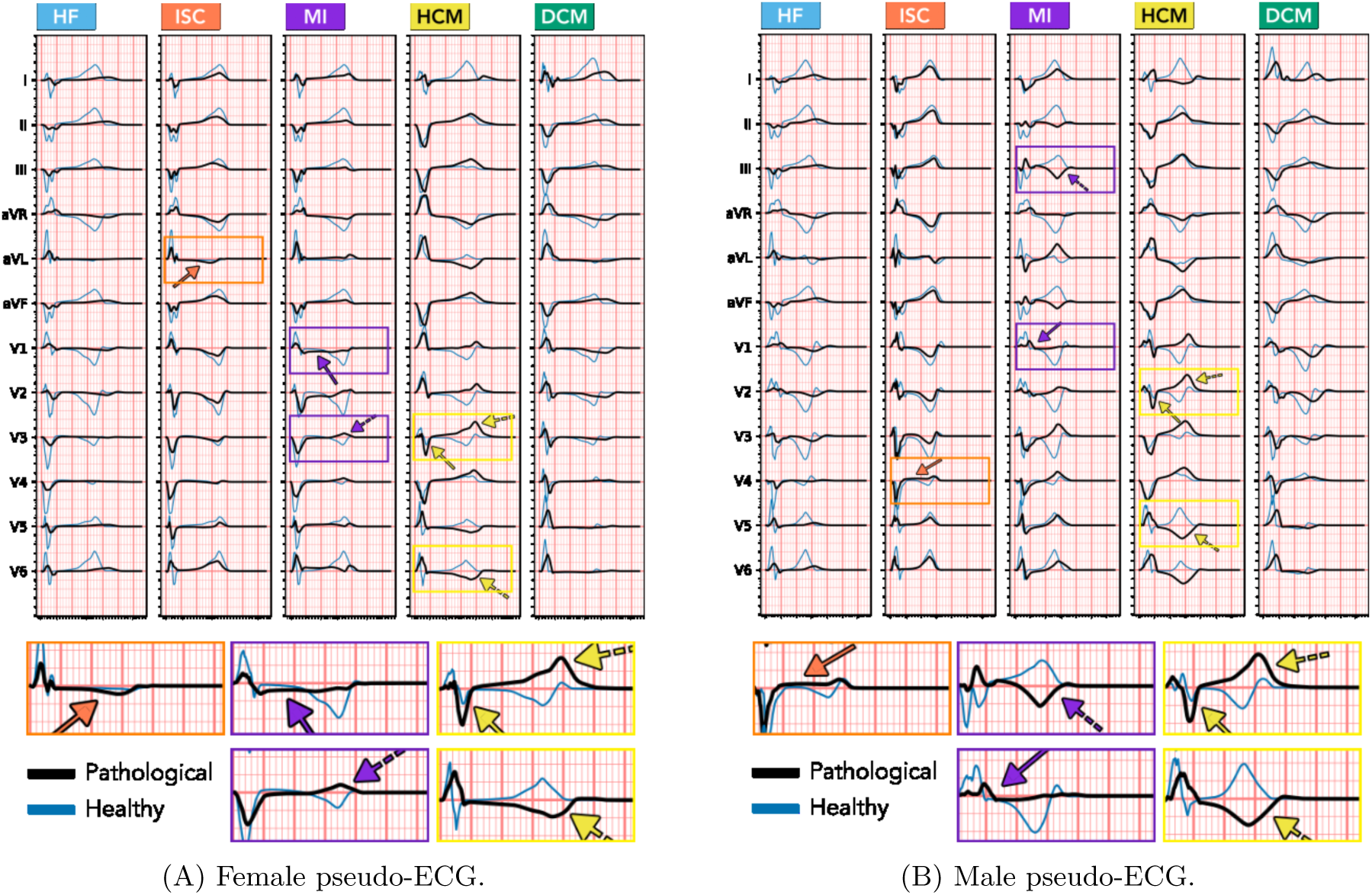
Examples of pseudo-ECG leads for a representative female (A) and male (B) case. Black lines correspond to pathological conditions, while blue lines represent the same leads under healthy conditions for the same subject. Yellow solid arrows highlight increased S-wave amplitudes; yellow dashed arrows indicate T-wave inversions or hyperacute T-waves. Orange solid arrows point to ST-segment depression in the female and ST-segment elevation in the male. Purple solid arrows denote ST-segment depression in the female and a pathological Q-wave in the male, while purple dashed arrows mark examples of T-wave inversions.

In HCM, characteristic ECG features include increased S-wave amplitude in leads V1-V3 and the presence of T-wave inversions or hyperacute T-waves, consistent with hallmark signs of the condition (Silvetti et al., 2023; Johnson et al., 2011). In ISC, sex-specific differences in myocardial scar type manifest as distinct ECG changes: ST-segment depression in females and ST-segment elevation in males, as reported by Gorgels (2013). Finally, in MI, scar type varies between sexes, resulting in ST-segment depression and T-wave inversions in females and pathological Q-waves with T-wave inversions in males, consistent with established infarction-related ECG patterns (Ferry, 2007; Higham et al., 1995).

In the healthy cohort, certain simulated ECGs exhibited atypical morphologies, including poor R-wave progression, inverted T-waves, and fragmented QRS complexes. These deviations are attributable to the simplified ventricular activation strategy adopted in the absence of an explicit Purkinje fiber network. Although such features introduce discrepancies in waveform morphology, they do not affect the interval-based biomarkers used for validation, which were consistent with reported physiological values, as explained in the following section.

In contrast, for local pathologies such as MI and ISC, the observed ECG changes were sometimes subtle (e.g., ST-segment deviations). This outcome is consistent with the clinical literature, as the surface ECG manifestations of these conditions are strongly determined by scar type and location.

### 3.2 Baseline Conditions

Figure 6 shows a violin plot for each anatomical group used to generate healthy and pathological populations, illustrating QT values at resting conditions. The violin plot displays the following metrics for QT values: the median is represented by a bold horizontal line in the centre of each box, while the mean is indicated by red dots along with their numerical values. The interquartile range is illustrated by the upper and lower lines. Within each range, the minimum and maximum values are represented by the length of the dispersion. Importantly, note that QT values are not directly correlated with the main quantity of interest, ΔQT, as described in Appendix C.

**Figure 6:**
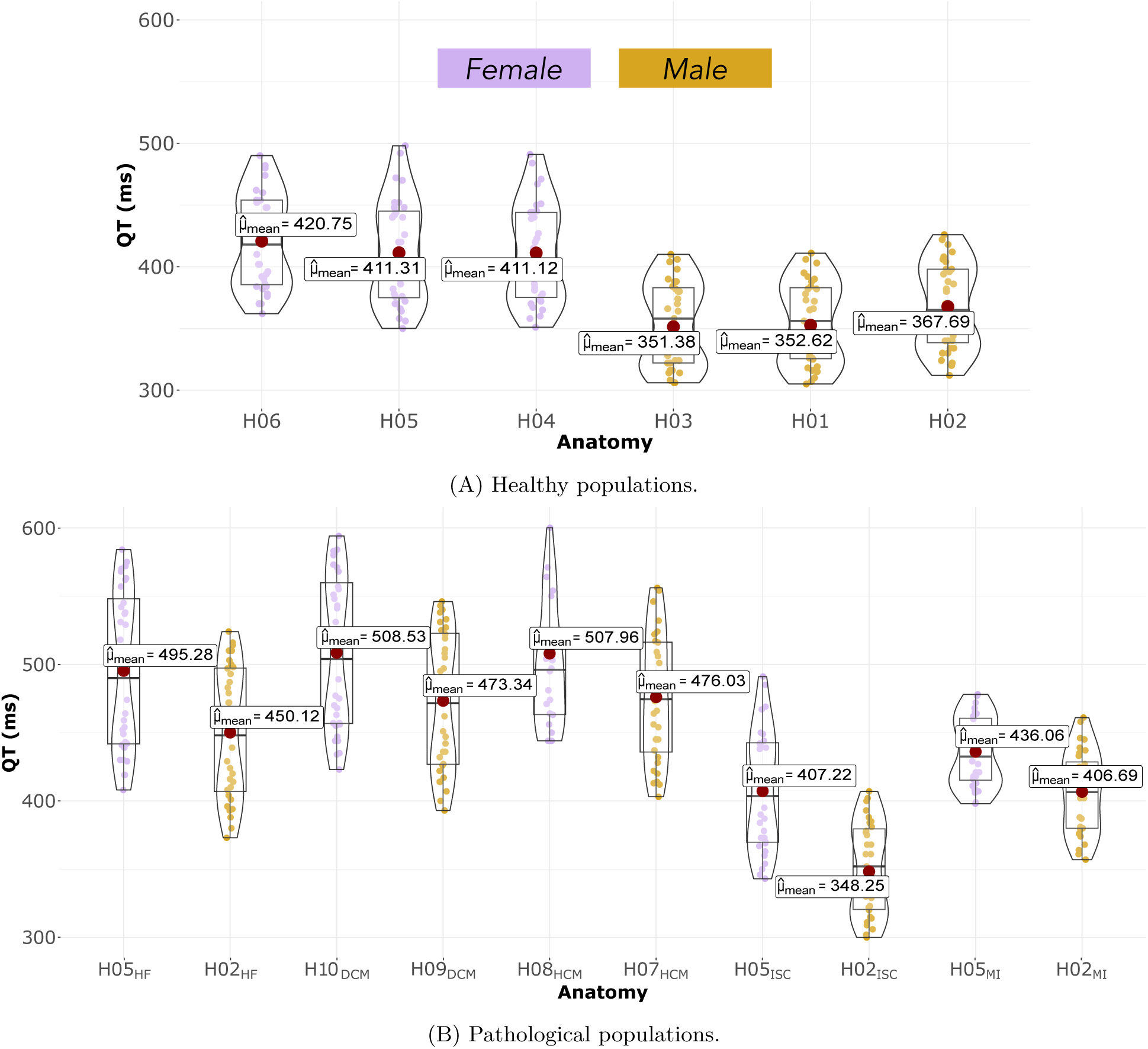
Comparison of QT values between healthy and pathological populations under baseline conditions.

Violin plots illustrate that healthy populations exhibit normal QT intervals, with females showing higher and more variable QT intervals compared to males, consistent with previous studies on sex-based electrophysiological differences (Darpo et al., 2014). In HF and DCM, QT intervals are elevated and more dispersed relative to healthy conditions, with females presenting higher QT values and a reduced gender gap in distribution, in agreement with reported data on QT prolongation in HF and DCM patients (Miró et al., 2023; Berger et al., 1997; Alonso et al., 2005).

In HCM, QT prolongation is generally observed only in a subset of patients (Johnson et al., 2011), largely due to the heterogeneous distribution of left ventricular hypertrophy, which has been shown to predispose to increased QT dispersion (Cava et al., 2023). However, in our HCM cohort, the distribution of hypertrophy in the whole left ventricle appears to result in a more generalized increase in QT intervals, which are also more prolonged and dispersed than in the healthy population. As in previous pathologies, females exhibited higher values, being eight excluded due to abnormal ECG patterns that made QT interval measurement impossible, a phenomenon consistent with arrhythmic prevalence in HCM (Tripathi et al., 2019).

In ISC, QT intervals remain within physiologically normal ranges, with females showing higher values. In MI, QT intervals also remain physiologically normal, although males exhibit slightly lower values and greater dispersion compared to females. Interestingly, two normal anatomies (female H05 and male H02) were employed to model HF, MI and ISC, so a direct comparison on the average effect of pathological states can be performed. ISC produced a 1% decrease of the average QT interval in the females and a 5% in the males. MI increased the QT interval by 6% in females, and by 9.6% in males. HF increased the average QT interval by 17% in females and 18.4% in males. The smaller differences observed in MI and ISC align with the reported clinical variability in comparison to other pathologies (Malik and Batchvarov, 2000; Higham et al., 1995). This smaller variability likely reflects the less extensive electrophysiological remodeling in MI and ISC, where scar burden was kept constant between sexes and the overall impact on repolarization is smaller than in chronic conditions like HF.

Table 8 presents a comparison between QT interval ranges obtained from clinical studies and those generated by our computational simulations. The simulated values demonstrate strong concordance with clinical data across all pathological conditions, thereby supporting the robustness of the computational methodology. For example, in the case of HF, reported clinical QT intervals range from 200 to 674 ms, whereas simulations produce values between 373 and 584 ms, effectively capturing the characteristic prolongation. In DCM, clinical QT intervals span 320 to 580 ms, while simulated intervals range from 393 to 594 ms—marginally higher but still consistent with the interindividual variability observed in patient populations.

**Table 8:**
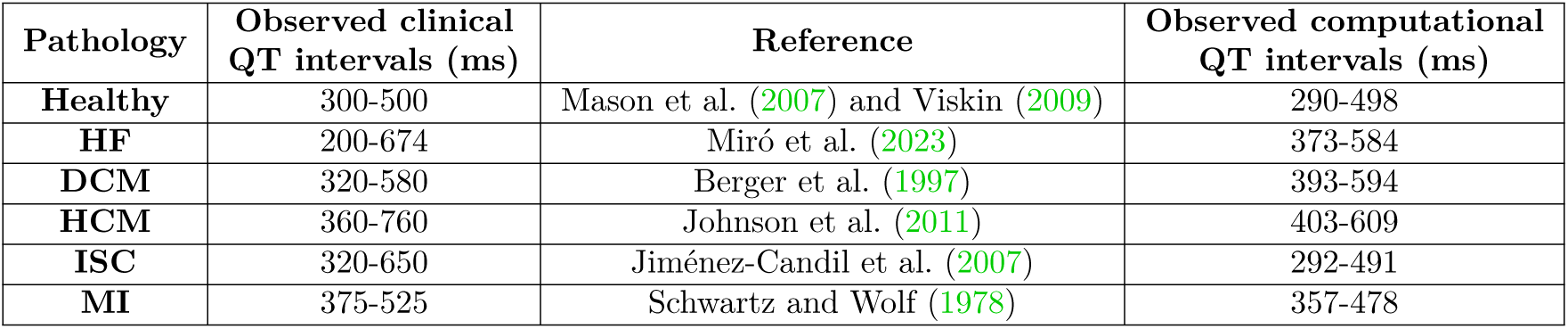
Observed QT interval ranges under different conditions in clinical practice and in the computational study.

In HCM, clinical QT intervals exhibit substantial variability (360–760 ms) reflecting the heterogeneous nature of hypertrophy. This variability is partially reproduced in the simulations, which yielded QT intervals ranging from 403 to 609ms, though extreme clinical values are not fully captured. For ISC and MI, both clinical and simulated QT intervals remain within physiological bounds, with simulations effectively representing the broader dispersion associated to ischaemia compared to infarction. Collectively, these results highlight the capability of computational models to replicate the complex relationship between pathology and repolarization abnormalities.

### 3.3 Benchmark Drugs

Figure 7 shows the concentration–response relationship for ΔQT across all conditions after the administration of moxifloxacin, while Table 9 summarizes the regression models’ statistics. In Appendix A (Figure A.9), violin plots illustrating ΔQT values by dose can be found.

**Figure 7:**
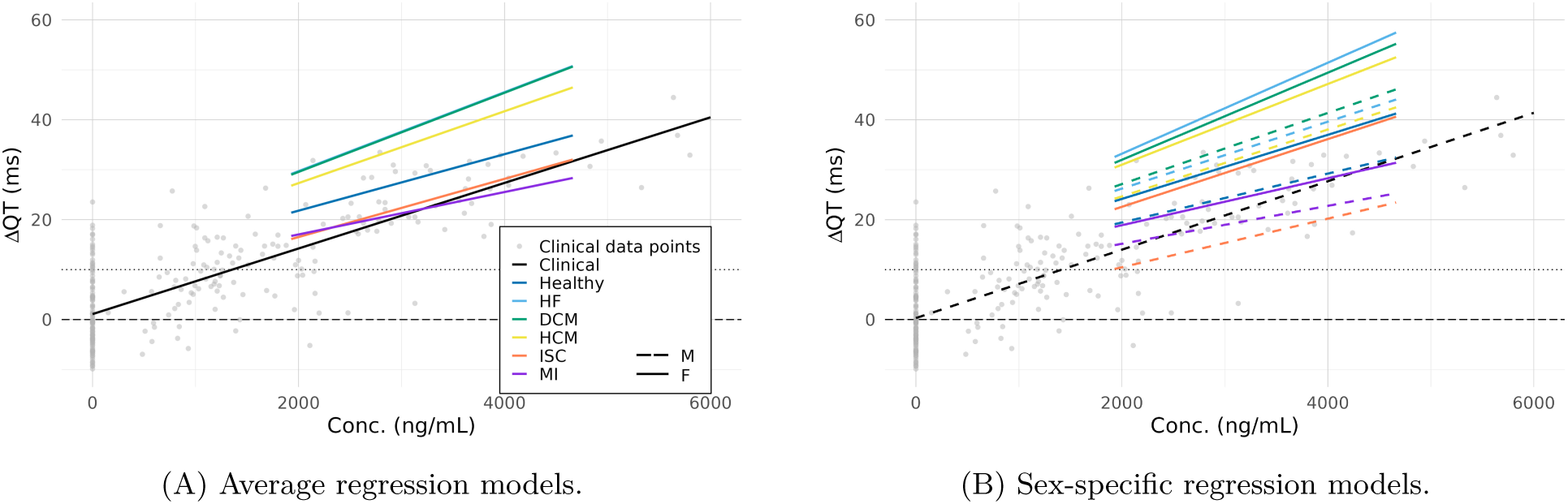
Concentration-QTc effect relationship models estimated for moxifloxacin from computational (healthy, HF, DCM, HCM, ISC and MI - ΔQT) and clinical data (healthy - placebo-adjusted ΔQT).

**Table 9:** Concentration-QTc effect relationship statistics estimated for moxifloxacin from computational (healthy, HF, DCM, HCM, ISC and MI - ΔQT) and clinical data (healthy - placebo-adjusted ΔQT). Key values are included, such as the slope, intercept, standard error of the slope, 90% confidence intervals, critical concentration, number of subjects, and the female-to-male ratio.

| Condition |  | Intercept | Slope | Slope std. error | Slope CI (Lower) | Slope CI (Upper) | Critical concentration | Total number of subjects | Female-to-male ratio (%) |
| --- | --- | --- | --- | --- | --- | --- | --- | --- | --- |
| Healthy | Avg | 10.485 | 0.006 | 0.0002 | 0.005 | 0.006 | - | 192 | 50 |
|  | F | 11.314 | 0.006 | 0.0004 | 0.006 | 0.007 | - |  |  |
|  | M | 9.657 | 0.005 | 0.0001 | 0.005 | 0.005 | - |  |  |
| HF | Avg | 13.910 | 0.008 | 0.0005 | 0.007 | 0.009 | - | 64 | 50 |
|  | F | 14.990 | 0.009 | 0.0007 | 0.008 | 0.010 | - |  |  |
|  | M | 12.829 | 0.007 | 0.0004 | 0.006 | 0.007 | - |  |  |
| DCM | Avg | 13.720 | 0.008 | 0.0005 | 0.007 | 0.009 | - | 64 | 50 |
|  | F | 14.552 | 0.009 | 0.0007 | 0.008 | 0.010 | - |  |  |
|  | M | 12.889 | 0.007 | 0.0005 | 0.006 | 0.008 | - |  |  |
| HCM | Avg | 12.920 | 0.007 | 0.0005 | 0.006 | 0.008 | - | 64 | 50 |
|  | F | 14.850 | 0.008 | 0.0009 | 0.007 | 0.010 | - |  |  |
|  | M | 11.267 | 0.007 | 0.0005 | 0.006 | 0.007 | - |  |  |
| ISC | Avg | 4.876 | 0.006 | 0.0006 | 0.005 | 0.007 | 879.938 | 64 | 50 |
|  | F | 9.019 | 0.007 | 0.0007 | 0.006 | 0.008 | 144.754 |  |  |
|  | M | 0.732 | 0.005 | 0.0002 | 0.005 | 0.005 | 1903.329 |  |  |
| MI | Avg | 8.543 | 0.004 | 0.0002 | 0.004 | 0.005 | 343.059 | 64 | 50 |
|  | F | 9.500 | 0.005 | 0.0001 | 0.004 | 0.005 | 106.333 |  |  |
|  | M | 7.586 | 0.004 | 0.0002 | 0.003 | 0.004 | 636.114 |  |  |
| Clinical | Avg | 1.094 | 0.007 | 0.0003 | 0.006 | 0.007 | 1356.450 | 9 | 0 |

In healthy subjects, moxifloxacin’s effect on ΔQT increases with dosage (400 ng/mL and 1700 ng/mL), with females showing higher and more dispersed ΔQT values than males. Mean ΔQT values (21.4 ms and 36.8 ms) slightly exceed the clinical benchmarks of 11.9 ms and 33.4 ms reported by Darpo et al. (2015), but still within their confidence interval, aligning with observations of QT prolongation trends across sexes.

Pathological conditions like HF, DCM and HCM exhibit elevated QT prolongation to moxifloxacin, as shown by higher intercepts and regression slopes, reflecting aggravated QT responses relative to the healthy population. Meanwhile, ISC and MI display relatively lower QT prolongation, in comparison to HF, DCM and HCM.

In all conditions, females consistently display slightly higher QT prolongation than males, what leads to regression analysis showing higher intercepts in female subjects, reflecting a lower critical concentration for QT response. Interestingly, the apparent widening of sex differences in ISC models does not arise from differences in ISC-induced changes themselves, which are the same for both sexes, but rather from the interaction of these changes with pre-existing structural differences due to scar distribution. Even if the sex-specific differences in scar characteristics are the same than those in MI models, the combination with ISC-induced electrophysiological changes produces a distinct effect on QT prolongation under the effect of moxifloxacin, amplifying the divergence between males and females.

### 3.4 Contraindicated Drugs

In the contraindicated drug analysis, each pathological condition was tested under specific drugs that are known or suspected to induce arrhythmic effects. No arrhythmia induction protocol was applied in this study. Instead, arrhythmias emerged spontaneously as a result of simulating each pathological condition under drug exposure. All ECG traces were visually inspected, and arrhythmias were manually classified according to the definitions summarized in Table 10.

**Table 10:** Number of individuals in the healthy and pathological populations according to the type of pseudo-ECG acquired: Normal (regular rhythm), PVC (Premature Ventricular Contraction), VT (Ventricular Tachycardia, a fast abnormal rhythm), TdP (Torsades de Pointes, a twisting VT), VF (Ventricular Fibrillation, chaotic rhythm), and Asystole (absence of electrical activity).

| Pathology | Drug | Sex | Dose 1 |  |  |  |  |  | Dose 2 |  |  |  |  |  |
| --- | --- | --- | --- | --- | --- | --- | --- | --- | --- | --- | --- | --- | --- | --- |
|  |  |  | Normal | PVC | VT | TdP | VF | Asystole | Normal | PVC | VT | TdP | VF | Asystole |
| HF | Quinidine | F | 28 | 2 | 2 | - | - | - | 6 | 2 | 6 | 5 | 9 | 4 |
|  |  | M | 32 | - | - | - | - | - | 22 | 8 | 2 | - | - | - |
| DCM | Bepridil | F | 32 | - | - | - | - | - | 26 | 5 | - | - | 1 | - |
|  |  | M | 32 | - | - | - | - | - | 32 | - | - | - | - | - |
| HCM | Bepridil | F | 19 | 8 | 5 | - | - | - | 16 | 6 | 4 | 2 | 4 | - |
|  |  | M | 32 | - | - | - | - | - | 32 | - | - | - | - | - |
| ISC | Flecainide | F | 32 | - | - | - | - | - | 20 | 5 | 1 | - | 6 | - |
|  |  | M | 32 | - | - | - | - | - | 32 | - | - | - | - | - |
| MI | Bepridil | F | 32 | - | - | - | - | - | 32 | - | - | - | - | - |
|  |  | M | 32 | - | - | - | - | - | 32 | - | - | - | - | - |

Results consistently highlighted that female patients showed greater susceptibility to arrhythmic behaviour, a trend observed across conditions and doses. These findings are summarized in Table 10.

For HF, quinidine results indicated significant QT prolongation, particularly among females, with a total of 30 abnormal ECG cases. At the second dose, 81% of the female subjects showed arrhythmias in comparison to 31% of male subjects. These results reveal a variety of arrhythmic patterns, including potentially lethal arrhythmias that were predominantly observed in female patients. Examples of these arrhythmic patterns can be found in Figure 8.

**Figure 8:**
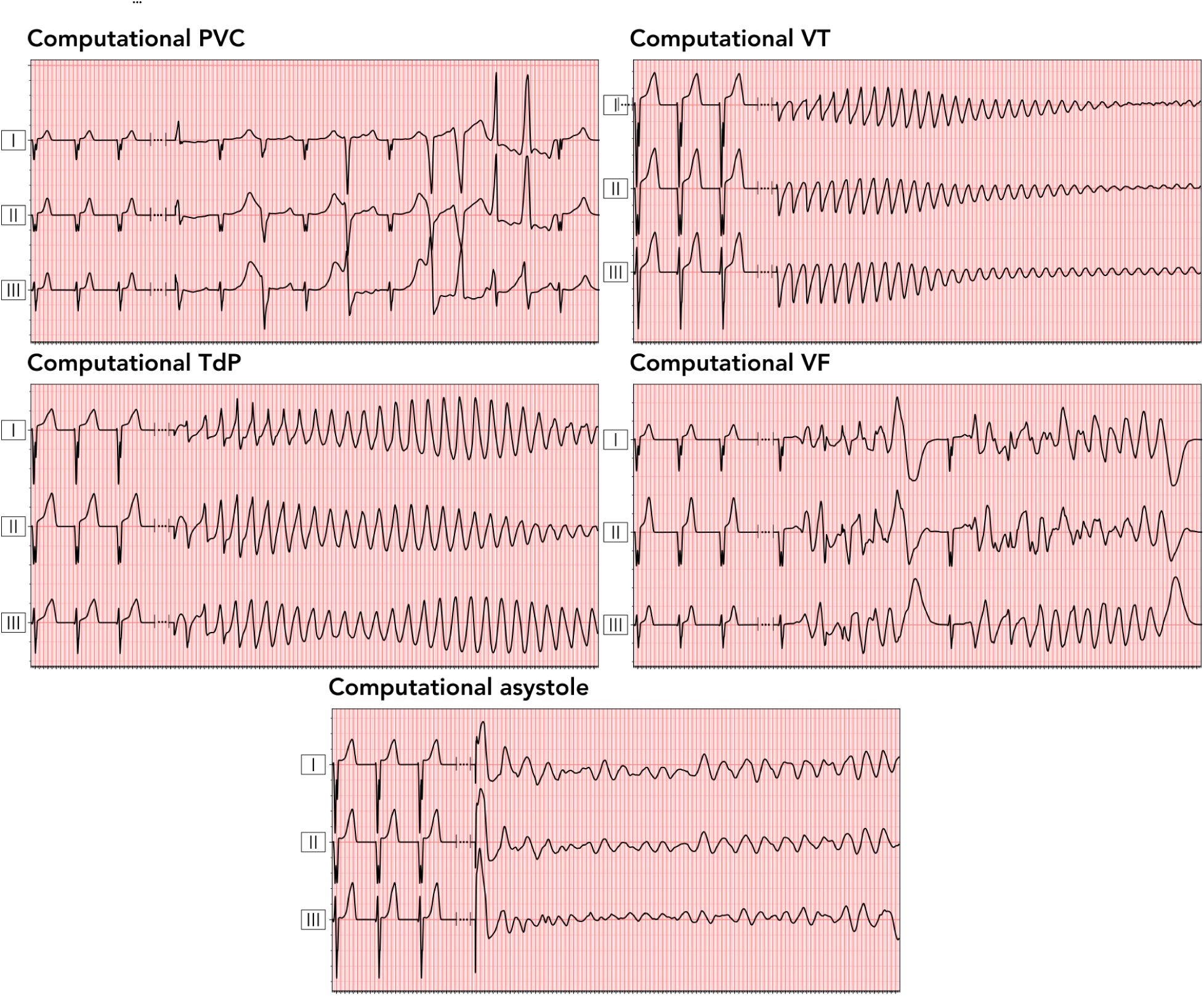
Computational pseudo-ECGs (leads I, II, and III) illustrating different arrhythmic events. Each trace begins with three stabilization beats under baseline conditions, followed—after a non-simulated transition period (indicated by a break)—by the ECG trace under drug exposure. This highlights the progression from normal rhythm to arrhythmia onset. Asystole trace shows very low amplitude, understood as residual noise below the threshold for effective ventricular activation.

DCM, HCM and MI populations were tested under the effect of bepridil. Female patients with DCM showed few arrhythmogenic cases, presenting only 19% of cases with abnormal ECG patterns at the second dose, with one being potentially lethal (VF); while males showed no arrhythmias. For HCM, however, bepridil’s effects were more pronounced, with a total of 29 cases of abnormal ECGs in females and none in males. Female subjects presented at least 6 lethal arrhythmias at the second dose. MI outcomes resulted in significant QT prolongation without arrhythmic outcomes in either sex. However, the second dose showed reversed T-wave inversion and caused ST segment elevation in four male cases.

In ISC, flecainide revealed arrhythmogenic effects on 37% of female subjects at the second dose, but none in males. Most importantly, 6 of those cases could be fatal.

The higher arrhythmic risk observed in HF compared with DCM, despite both using the same electrophysiological remodeling, can be partly explained by the different anatomical substrates and the drugs used for testing. While DCM anatomies were artificially dilated, with larger chambers and thinner walls that might be expected to promote reentry, the HF population was tested with quinidine, while DCM was tested with bepridil, whose mixed ion-channel block produces a milder proarrhythmic response. The combination of HF geometry with quinidine’s pronounced effect on repolarization likely enhanced susceptibility to torsadogenic events, leading to the higher arrhythmic burden.

Individuals who developed arrhythmias across the different pathological populations exhibit similar phenotypic expression profiles. This suggests that the combination of sex-specific electrophysiological traits and interindividual variability creates a substrate prone to arrhythmogenesis, with specific disease conditions acting as triggers. Notably, susceptible individuals tend to share a common ionic remodelling pattern—characterised by stronger block of *I*_Kr_ and *I*_Ks_, and increased *I*_CaL_ conductance—which amplifies repolarization abnormalities and facilitates arrhythmic events across pathologies.

These arrhythmogenic effects were accompanied by substantial QT prolongation across all tested conditions for the non-arrhythmic cases, as summarized in Table 11. In line with previous findings, females exhibited a consistently greater increase in QT interval compared to males. The most pronounced prolongations were observed in HF patients treated with quinidine, where QT values reached 329 ms in females and 266 ms in males after the second dose. Similarly, bepridil produced notable QT prolongation in DCM, HCM, and MI populations, with the highest values in females with DCM (149 ms after the second dose). Flecainide also resulted in a significant QT increase in ISC, particularly in females (161 ms).

**Table 11:** Mean QT interval values for non-arrhythmic cases in the pathological populations under the effect of the contraindicated drugs. N accounts for the number of subjects.

| Pathology | Drug | Sex | Dose 1 |  | Dose 2 |  |
| --- | --- | --- | --- | --- | --- | --- |
| | | | N | Mean $\Delta QT$ (ms) | N | Mean $\Delta QT$ (ms) |
| HF | Quinidine | F | 28 | 119 | 6 | 329 |
|  |  | M | 32 | 94 | 22 | 286 |
| DCM | Bepridil | F | 32 | 66 | 26 | 149 |
|  |  | M | 32 | 56 | 32 | 128 |
| HCM | Bepridil | F | 19 | 58 | 16 | 142 |
|  |  | M | 32 | 48 | 32 | 106 |
| ISC | Flecainide | F | 32 | 64 | 20 | 161 |
|  |  | M | 32 | 42 | 32 | 120 |
| MI | Bepridil | F | 32 | 36 | 32 | 84 |
|  |  | M | 32 | 33 | 32 | 71 |

## 4 Discussion

This work illustrates the utility of virtual populations and underscores the important role of computational modelling in evaluating drug-induced proarrhythmic risk across a wide spectrum of patient profiles. This includes populations frequently underrepresented in cardiac safety clinical trials, such as females and patients with preexisting heart conditions. Through the creation of virtual cohorts for both healthy and pathological populations, this study provides critical insights into the differences in response to QT-prolonging drugs, highlighting sex-specific as well as condition-dependent variability and uncovering nuanced arrhythmic risks that may otherwise be overlooked in cardiac safety clinical trials.

Female subjects consistently showed higher QT and ΔQT values, along with greater variability, across baseline conditions and under the effects of moxifloxacin, as illustrated in Figures 6 and 7. Higher QT and ΔQT values in females align with clinical findings (Darpo et al., 2014) and highlight that sex-specific variability, often limited by an overrepresentation of male participants in clinical trials, could influence QT prolongation outcomes. These findings support the need for sex-balanced cohorts in drug safety research. Computational modelling that captures these variations offers a more nuanced view of QT prolongation risks, addressing critical gaps left by cardiac safety clinical trials that potentially underestimate drug-related risks for female. Differences in sample size and sex ratios highly affect statistical outcomes in clinical trials (Reza et al., 2022; Melloni et al., 2010). The balanced male-to-female ratio in this computational study contrasts with male-dominated clinical trials, highlighting potential biases in reported QT prolongation that could impact risk stratification and drug safety profiles.

Beyond sex differences, our analysis reveals distinct drug responses across pathological conditions. Conditions such as HF, HCM, and DCM demonstrated higher sensitivity to potentially QT-prolonging drugs, as observed for moxifloxacin (Figure 7 and Table 9) but presented no arrhythmias overall. ISC and MI presented lower overall QT prolongation. This condition-specific response to QT-prolonging drugs highlights the heterogeneous nature of cardiac pathologies and emphasizes the importance of tailoring arrhythmia risk assessments to the individual’s underlying heart condition. The differential response in HF versus ISC patients suggests that pharmacological strategies may need to be assessed in a condition-specific manner, with targeted risk management protocols to address each condition’s unique arrhythmic profile.

Another key observation from this study is that ISC conditions exhibited a greater divergence between male and female responses compared with other pathological conditions, particularly MI (Table 9). Even though the male–female difference in scar characteristics was the same as in the MI models, the combined effect of ischemic remodeling and scar in ISC resulted in a more pronounced separation in QT responses. This widening of sex differences under ISC likely reflects a synergistic interaction between ischemic remodeling and scar structural heterogeneities that is exhacerbated by the effect of moxifloxacin.

Arrhythmic behaviours observed in the study, including PVCs, VT, TdP, VF, and Asystole, were more prevalent among female subjects across several pathological groups, especially in response to contraindicated drugs. Table 10 summarizes these arrhythmic outcomes, showing quinidine’s significant QT-prolonging and arrhythmic effects in HF patients, flecainide’s mild arrhythmogenic effects in ISC patients, and bepridil’s differential effects on DCM and HCM patients, which induced 6 and 29 cases of abnormal ECGs in females compared to none in males. This result aligns with studies suggesting that female HCM patients may have a poorer prognosis, highlighting a potential gender-based differential response in arrhythmogenic risk (Butters et al., 2021). MI outcomes resulted in significant QT prolongation without arrhythmic outcomes in either sex. However, in four male cases, the second dose reversed T-wave inversion and caused ST segment elevation, suggesting a possible protective effect that aligns with studies noting bepridil’s antiarrhythmic role in post-infarct conditions (Flammang et al., 1989; Lynch et al., 1986). These findings suggest that the use of computational models that incorporate sex and pathological variability can serve as predictive tools for identifying at-risk populations for drug-induced arrhythmia. Furthermore, it was able to provide information of potentially lethal events, providing an early stratification mechanism for the use of common drugs on vulnerable populations. Notably, the common phenotype among arrhythmia-prone individuals—characterised by stronger *I*_Kr_ and *I*_Ks_ block and enhanced *I*_CaL_ —is consistent with the well-established role of *I*_Kr_ as a key determinant of delayed repolarization and drug-induced proarrhythmia (Mitcheson et al., 2000), reinforcing the physiological relevance of our findings.

The analysis of contraindicated drugs further underscores the well-documented sex differences in drug-induced QT prolongation that magnify as the risk increases (Darpo et al., 2014), clearly enhanced by the presence of some pathologies, such as HF (Table 11). Our findings show that, the combined effect of disease and drug exposure leads to extreme QT prolongation and a heightened likelihood of arrhythmic events, further exacerbating the known sex disparities. This suggests that underlying disease states not only modulate the extent of QT prolongation but also amplify pre-existing sex differences, potentially explaining why female patients experience a greater incidence of drug-induced arrhythmias in clinical settings. These insights further emphasize the need for sex-specific considerations in pharmacological safety assessments, particularly in populations with cardiac comorbidities.

Due to the currently limited availability of clinical data on drug-induced cardiac safety in pathological populations, direct validation of computational predictions often becomes feasible only upon the emergence of clinical arrhythmic events. This context highlights the importance of continued efforts to expand and curate high-quality datasets. In this study, the inclusion of contraindicated drugs served as a strategic “negative control”, enabling the assessment of potential arrhythmic risks beyond standard treatments. Furthermore, increasing the diversity and variability within virtual patient populations represents a promising direction for enhancing the physiological realism of in silico trials and improving the generalizability of model-based predictions to real-world clinical settings.

Although this study provides valuable insights, it has some limitations. The pathological models could be refined further to capture more complex biological mechanisms associated to the selected diseases. The current models are limited by the lack of cardiac contraction, which constrains the full replication of the dysfunctions typically seen in real-world cardiac pathologies. Additionally, genetic-based diseases or those involving specific mutations were not the target of this study, potentially overlooking critical disease characteristics. While enhancing model complexity could address these limitations, it would also increase computational demands.

Another limitation relates to ECG morphology. Due to the simplified ventricular activation sequence used in the absence of a detailed Purkinje conduction system, some simulated ECGs displayed atypical features such as poor R-wave progression, inverted T-waves, or fragmented QRS complexes. These morphological discrepancies do not reflect clinical healthy ECGs but are a known artifact of whole-heart simulations without explicit Purkinje networks. Importantly, they did not affect the interval-based biomarkers employed for validation, which remained within reported physiological ranges.

Furthermore, the electrophysiological models used in this study are based on the ORd model. While this approach has been widely used, more recent models, such as the TorORD model (Tomek et al., 2019), have been developed since 2016. Future studies could explore the implications of incorporating newer models to assess potential differences in drug response predictions.

Population variability was introduced by independently scaling the conductances of five major ionic currents (*I*_Na_, *I*_Kr_, *I*_Ks_, *I*_CaL_, and *I*_NaL_). While this captures much of the action potential variability observed experimentally, it omits potential variability in other currents and parameters that may influence repolarization and arrhythmic susceptibility. Expanding the parameter space in future work could improve physiological coverage. Moreover, while our study assumes independent variability in ion channel conductances, experimental evidence suggests that some ionic currents are coregulated (Ballouz et al., 2020), particularly under pathological or hormonal influences. However, capturing such relationships in computational models remains challenging due to the lack of mechanistic understanding and robust quantitative data needed to define such relationships in a generalizable way. Our model also assumes homogeneous distributions of ion channel conductances across endo-, mid-, and epicardial cells, thereby omitting known transmural heterogeneities. Future work could aim to incorporate both coregulatory patterns and transmural variability to more accurately reflect the full spectrum of electrophysiological diversity observed in human hearts.

Finally, the IC50 values and Hill coefficients used in this study were derived from a single dataset, despite the wide range of values reported in the literature. Future studies should consider integrating multiple datasets or exploring uncertainty quantification techniques to assess the impact of variability in drug-binding parameters on model predictions.

### 4.1 Conclusions

This study addresses a critical challenge in cardiac drug safety: the underrepresentation of high-risk populations in traditional proarrhythmic risk assessments. By employing cardiac computational models of both healthy and pathological populations, we demonstrate that in silico trials can reveal nuanced drug-induced responses that may remain undetected in conventional clinical studies.

Our findings show that female virtual subjects consistently exhibit higher baseline QT values and greater sensitivity to drug-induced QT prolongation, both in healthy and pathological conditions. The use of moxifloxacin and contraindicated drugs enabled us to explore a range of proarrhythmic risks across healthy, HF, DCM, HCM, ISC and MI populations. While moxifloxacin led to significant QT prolongation—particularly in HF, DCM, and HCM—only the contraindicated drug triggered severe arrhythmic events such as TdP, VF, and asystole, predominantly in female subjects. These results highlight condition- and sex-specific responses to QT-prolonging agents and support the utility of computational models in early risk stratification.

By capturing both sex- and disease-related variability, this work lays the foundation for a framework to proactively assess arrhythmic risk in vulnerable populations, supporting more inclusive and, therefore, reliable safety evaluations. It addresses important gaps in current clinical practices and highlights the growing potential for in silico trials to become key components of preclinical decision-making. As models continue to improve, their integration into drug development pipelines could significantly help mitigate unforeseen risks, guide safer clinical trial design, and promote more equitable pharmacological innovation.

## 5 Additional Information

### Author Contributions

We would like to acknowledge the significant contributions made by all authors to this work. P.D.-G. and P.G.-M. contributed to the conception and design of the work, data curation, formal analysis, investigation, methodology, project administration, resources, validation, visualization, writing - original draft, and writing

– review & editing. L.B.-C., E.C. and A.A. contributed to the conception and design of the work, data curation, formal analysis, investigation, methodology, resources, software, writing - original draft, and writing
– review & editing. J.M.P. and C.B. contributed to the conception and design of the work, methodology, resources, software, writing - original draft, and writing - review & editing. M.V. contributed to funding acquisition, project administration, resources, software, supervision, and writing - review & editing. J.A.-S. contributed to the conception and design of the work, data curation, formal analysis, funding acquisition, investigation, methodology, project administration, resources, software, supervision, validation, visualization, writing - original draft, and writing - review & editing.

All authors approved the final version of the manuscript and agree to be accountable for all aspects of the work, ensuring that any questions related to the accuracy or integrity of the work are appropriately investigated and resolved. All authors meet the criteria for authorship and are listed accordingly.

## Data Availability

Elem Biotech owns the commercial rights to Alya, the computational finite element solver employed in this study for the simulator. The methodology can be replicated using any finite element solver given all the models’ information provided in this paper.

The data generated and analysed during this study are proprietary to Elem Biotech and cannot be publicly shared due to commercial confidentiality. However, access to the data may be considered on a case-by-case basis. Requests for access can be directed to Elem Biotech compliance department, and will be subject to a data use agreement ensuring compliance with relevant regulations and restrictions.

## Appendix A

### Benchmark Drugs

Figure A.9 depicts the ΔQT values for both doses of moxifloxacin across all virtual populations, highlighting the drug’s dose-dependent effects on QT prolongation and variability across healthy and pathological cohorts.

**Figure A.9:**
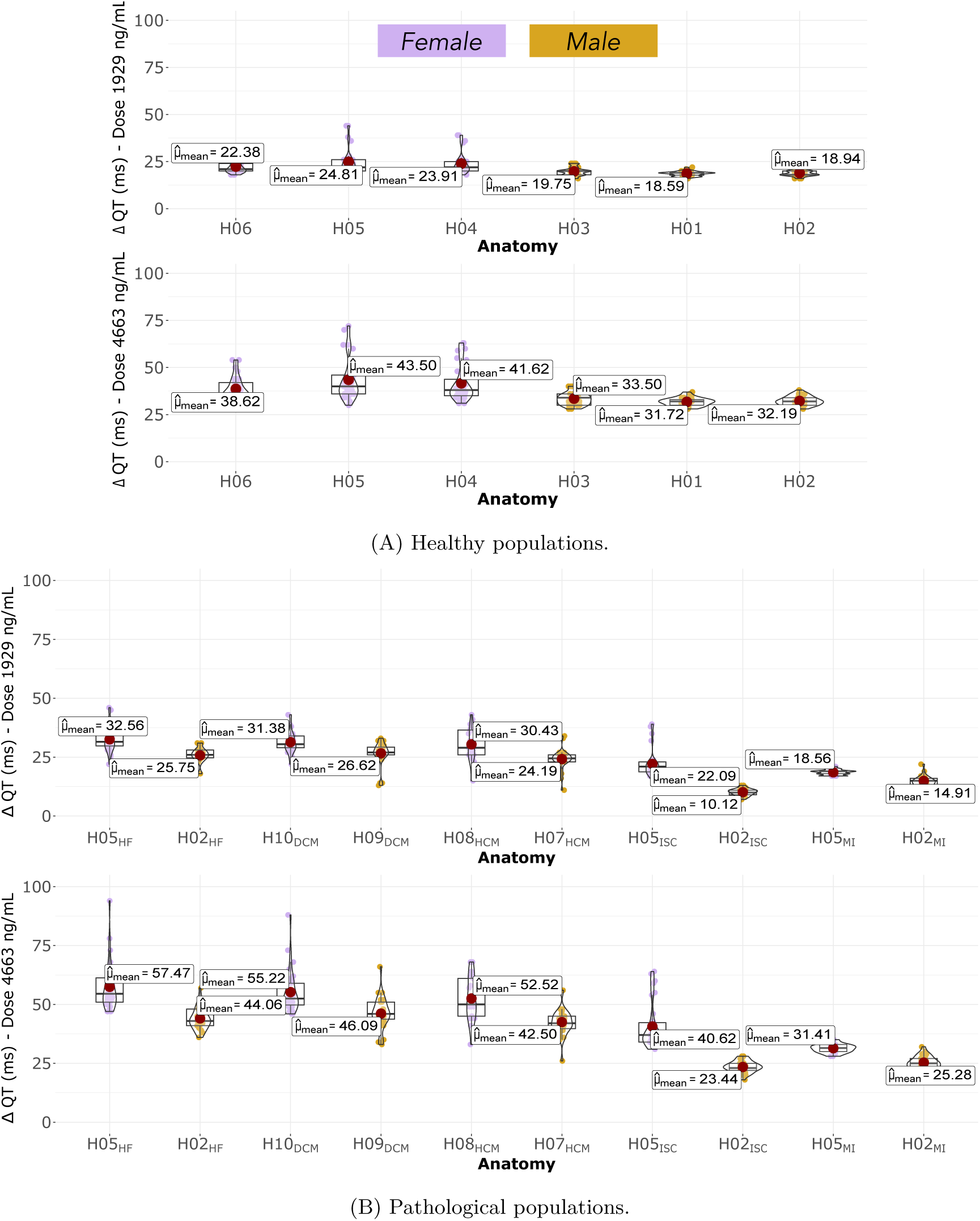
Comparison of ΔQT values between healthy and pathological populations subjected to the effect of moxifloxacin.

## Appendix B

### Artificial Detailed DCM hearts

The detailed DCM hearts used in this study were generated by morphing a high-resolution healthy heart model into low-resolution in vivo-based DCM hearts. The in vivo DCM hearts were reconstructed from cine magnetic resonance imaging (MRI) of patients with left bundle branch block. Two DCM hearts were selected based on their left ventricular end-diastolic volume (LVEDV), normalized to body surface area (BSA), measuring 237.35 *ml/m*^2^ and 263.23 *ml/m*^2^, respectively. Both values exceeded twice the normal standard deviation (Z-score > 2) (Pinto et al., 2016).

The healthy heart model, H05, was morphed into the selected in vivo DCM hearts using an in-house morphing algorithm to create the artificially detailed DCM hearts. In this process, the in vivo DCM heart served as the reference (target) mesh, while the high-resolution H05 heart model was morphed to match the target mesh as closely as possible. A schematic representation of this process is shown in Figure B.10.

**Figure B.10:**
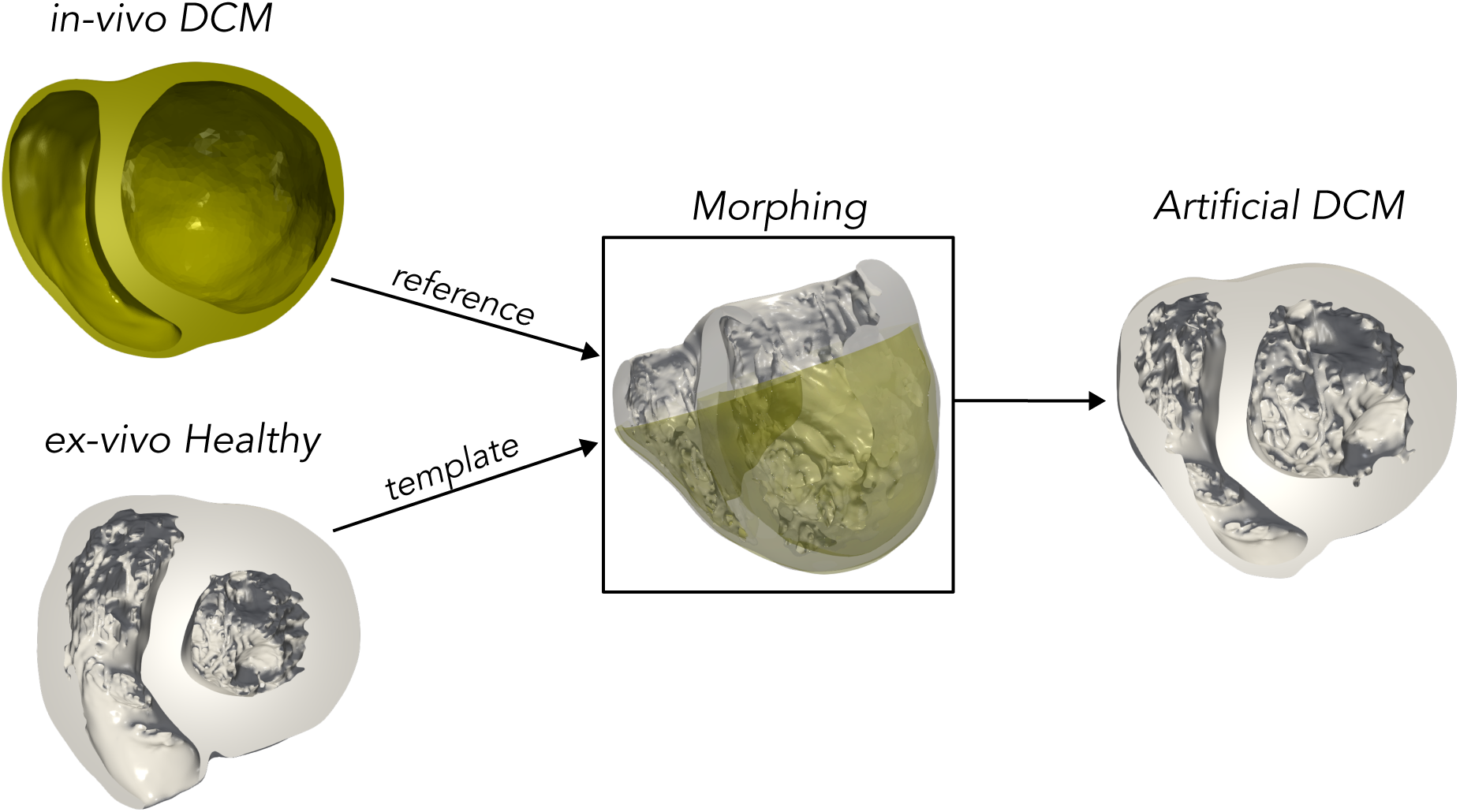
Schematic representation of the generation of artificial DCM cases.

## Appendix C

### Non-Dependence of Drug-Induced **Δ**QT responses on QT Values

As shown in Figure 6, some male and female models exhibit QT values near or slightly outside the expected healthy QT ranges. These extremes arise from the physiological variability embedded in our population modeling framework and are within the expected dispersion given the stochastic parameter sampling. However, we emphasize that the presence of such extreme QT values does not compromise the interpretation of our main quantity of interest, ΔQT.

To assess whether these unusually low (or high) baseline QT intervals influence ΔQT results, we conducted an additional set of simulations. Specifically, we selected a male healthy baseline population with QT intervals at the lower extreme, and generated a population variant adjusted to more closely match the clinical average. As shown in Figure C.11, the resulting ΔQT values are consistent with those obtained from the original population, indicating no strong dependence of ΔQT on absolute QT interval duration.

**Figure C.11:**
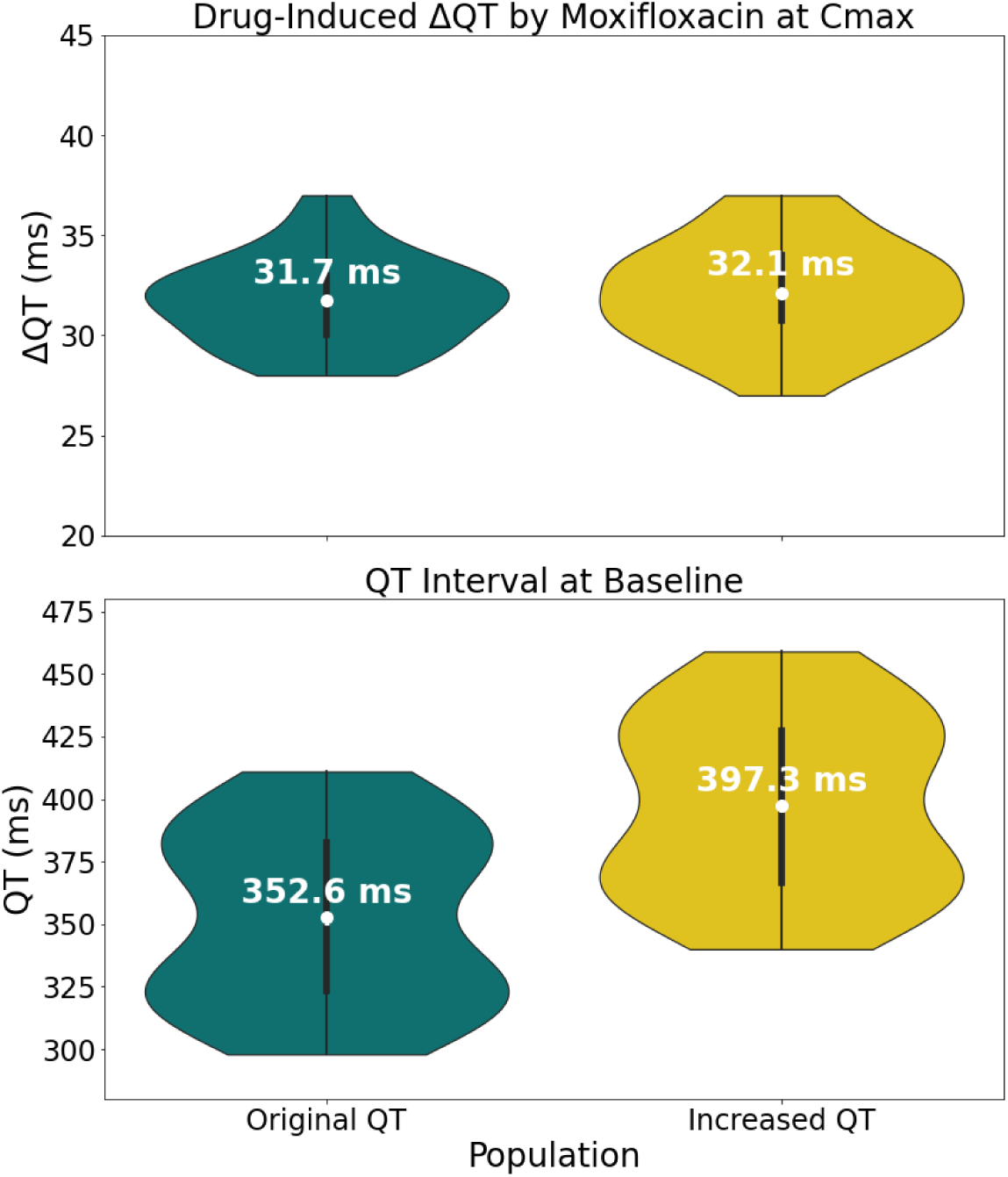
ΔQT values under the effect of Moxifloxacin at Cmax in a male population with different QT intervals under baseline conditions.

